# Cortical beta oscillatory activity evoked during reactive balance recovery scales with perturbation difficulty and individual balance ability

**DOI:** 10.1101/2020.10.07.330381

**Authors:** Nina J. Ghosn, Jacqueline A. Palmer, Michael R. Borich, Lena H. Ting, Aiden M. Payne

**Author notes:** **Corresponding Author:** Dr. Aiden Payne, PhD Postdoctoral Research Fellow Biomedical Engineering, Emory University School of Medicine 1760 Haygood Dr NE, Ste W200, Atlanta, GA 30322.

## Abstract

Cortical beta oscillations (13-30 Hz) reflect sensorimotor cortical activity, but have not been fully investigated in balance recovery behavior. We hypothesized that more challenging balance conditions would lead to greater recruitment of cortical sensorimotor brain regions for balance recovery. We predicted that beta power would be enhanced when balance recovery is more challenging, either due to more difficult perturbations or due to lower intrinsic balance ability. In 19 young adults, we measured beta power evoked over motor cortical areas (Cz electrode) during 3 magnitudes of backward support-surface translational perturbations using electroencephalography. Peak beta power was measured during early (50-150 ms), late (150-250 ms), and overall (0-400 ms) time bins, and wavelet-based analyses quantified the time course of evoked beta power and agonist and antagonist ankle muscle activity. We further assessed the relationship between individual balance ability measured in a challenging beam walking task and perturbation-evoked beta power within each time bin. In balance perturbations, cortical beta power increased ∼50 ms after perturbation onset, demonstrating greater increases with increasing perturbation magnitude. Balance ability was negatively associated with peak beta power in only the late (150-250 ms) time bin, with higher beta power in individuals who performed worse in the beam walking task. Additionally, the time course of cortical beta power followed a similar waveform as the evoked muscle activity, suggesting these evoked responses may be initially evoked by shared underlying mechanisms. These findings support the active role of sensorimotor cortex in balance recovery behavior, with greater recruitment of cortical resources under more challenging balance conditions. Cortical beta power may therefore provide a biomarker for engagement of sensorimotor cortical resources during reactive balance recovery and reflect the individual level of balance challenge.

## II. Introduction

In response to a destabilizing event, standing balance-correcting behavior is comprised of a consistent, fast, and automatic brainstem-mediated response (Deliagina et al., 2006, 2007; Horak & Macpherson, 1996; Welch & Ting, 2008, 2009) followed by a more variable later motor response (Jacobs & Horak, 2007; B. E. Maki & McIlroy, 2007). Due to the longer latency of its occurrence (>150 ms post perturbation onset), it has been postulated that the later motor component of the balance reaction is actively influenced by inputs from higher brain centers, including the cerebral cortex. This is further supported by the presence of severe balance impairments in humans with cortical damage, such as after cortical stroke (Lin et al., 2014; Patel & Bhatt, 2016; Pérennou et al., 2000), and the deterioration of reactive balance function that occurs when the cortex is engaged in simultaneous performance of a cognitive dual task (Bolton et al., 2011; Jacobs & Horak, 2007; B. E. Maki & McIlroy, 2007). Behavioral studies involving cognitive dual task paradigms also implicate greater cortical contributions to balance control in individuals with lower balance ability (Jacobs & Horak, 2007; B. E. Maki & McIlroy, 2007; Shumway-Cook et al., 1997), particularly in aging adult populations (Rankin et al., 2000). However, there have been limited direct investigations of the cortical mechanisms involved in balance control (Stuart et al., 2018). We previously characterized the negative cortical event-related potential (N1) response during balance reactions in young adults using electroencephalography (EEG). We found that evoked cortical N1 responses, thought to reflect general attention (C. E. Little & Woollacott, 2015; Quant et al., 2004) and perceived threat (Adkin et al., 2008; Payne et al., 2019), were larger in young adults with lower balance ability (Payne & Ting, 2020). As a follow-up to the prior study, we now aim to characterize signatures of *sensorimotor information processing* during balance recovery by quantifying cortical activity in the beta frequency range (13-30 Hz) throughout balance perturbations at different levels of perturbation magnitude and in individuals with different levels of balance ability.

Oscillatory activity in the beta frequency range is a prevalent feature of sensorimotor neural networks and is closely linked to motor behavior. While beta oscillations are robustly suppressed at the initiation of voluntary reaching behaviors (van Wijk et al., 2012; Zaepffel et al., 2013), beta oscillatory activity is enhanced and sustained during isometric motor activity when maintaining a fixed arm posture (van Wijk et al., 2012; Zaepffel et al., 2013). Direct modulation of cortical beta activity using transcranial alternating current stimulation delivered at beta frequency ranges can slow (Joundi et al., 2012; Pogosyan et al., 2009) or reduce the amplitude (Guerra et al., 2018) of ongoing motor activity, providing evidence for the causal role of cortical beta oscillations in sustained ongoing motor behavior.

One plausible theory for beta power modulation during sustained motor activity is that beta oscillations reflect cortical network activity contributing to maintaining the “status quo” of a given motor state (Engel & Fries, 2010). Cortical beta oscillations appear to reflect inhibitory γ-aminobutyric acid (GABA) neural network activity (Stuart N. Baker, 2007; Muthukumaraswamy et al., 2013). Abnormal movement-related beta oscillatory modulation and GABAergic activity have been well-established in elderly adults and other neurologic populations with impaired balance (Heise et al., 2013; H. E. Rossiter et al., 2014; Holly E. Rossiter et al., 2014). In these patient populations, beta oscillatory activity has been linked to impaired volitional motor activity (Rossiter et al., 2014a; Rossiter et al., 2014b) and impaired ability to “switch” from the current motor state, e.g. difficulty initiating volitional movements or halting ongoing ambulation in Parkinson’s disease (Brown, 2003; Kühn et al., 2006, 2009). This evidence raises the possibility that measures of beta oscillatory activity may provide useful insight into sensorimotor cortical mechanisms involved in postural control and balance responses.

While beta oscillations have been linked to sensorimotor information processing and motor control, beta oscillatory activity in the context of balance-correcting behavior and associations with balance ability are poorly understood. Previous studies have consistently demonstrated that cortical beta activity over central midline scalp regions is modulated during standing balance recovery (Nakamura et al., 2020; Peterson & Ferris, 2018, 2019; Solis-Escalante et al., 2019; Varghese et al., 2014, 2019), though there is some discrepancy regarding the directionality of change that may originate from differences in the perturbation and analysis methodologies.

Although balance perturbations evoke a robust modulation of cortical beta activity, the *precise time course* of modulation remains unclear, limiting our understanding of potential causal contributions of sensorimotor cortical activity to reactive balance control. Further, it is conceivable that higher level cortical brain regions may only be recruited when balance is considerably challenged, when automatic brainstem-mediated balance reactions are insufficient for balance recovery. This notion is supported by cognitive dual task studies showing that the greatest effect of cognitive interference occurs during the most difficult balance conditions (Lajoie et al., 1993; Woollacott & Shumway-Cook, 2002) and in individuals with worse balance (Shumway-Cook et al., 1997). However, the variations in cortical beta activity with balance task difficulty and individual differences in balance ability have not been investigated previously and could help elucidate the role of sensorimotor cortical involvement in balance control.

Here we hypothesized that engagement of sensorimotor cortical brain regions would increase with the level of individual balance difficulty. As such, we predicted that modulation of cortical beta activity during reactive balance recovery would be dependent on both perturbation magnitude and individual balance ability. In young adults, we aimed to 1) determine the within-individual effect of balance perturbation magnitude on beta activity modulation, 2) test the between-individuals relationship between cortical beta activity during reactive balance recovery and balance ability as measured in a challenging beam walking task, and 3) characterize the precise time course of beta activity modulation during reactive balance recovery with respect to the evoked muscle activity using wavelet decomposition analyses. We predicted that the modulation of cortical beta activity would scale with balance perturbation magnitude, with the greatest cortical beta activity modulation occurring at the highest level of perturbation magnitude where cortical resources are more likely to be recruited for balance recovery. We expected that individuals with the lowest balance ability would show the greatest modulation of cortical beta activity, since they should experience a relatively greater challenge at a given level of perturbation magnitude, necessitating greater recruitment of sensorimotor cortical resources for successful balance recovery.

## III. Methods

### Participants

Nineteen young adults (11 female, ages 19-38 years) were recruited from Emory University and the surrounding Atlanta community to participate in a single experimental testing session. Exclusion criteria included history of neurologic or musculoskeletal pathology or vision impairment that could not be improved with corrective lenses to 20/40 or better. Different analyses of this same participant cohort have been reported previously (Payne & Ting, 2020). Participants were 26±5 years old (mean ± standard deviation), 168±8 cm tall, and 70±14 kg. The experimental protocol was approved by the Emory University Institutional Review Board and all participants provided written informed consent in accordance with the Declaration of Helsinki.

### Balance Perturbations

While participants stood quietly and barefoot on a custom platform (Factory Automation Systems, Atlanta, GA), 48 backward translational support-surface perturbations were delivered at unpredictable timing and in randomized order of magnitude (i.e. difficulty) (Payne & Ting, 2020). Participants received an equal number of perturbations in each of three magnitudes: a small perturbation (7.7 cm, 16.0 cm/s, 0.23 g) that was identical across participants, and two larger magnitudes (medium 12.6-15.0 cm, 26.6-31.5 cm/s, 0.38-0.45 g and large 18.4-21.9 cm, 38.7-42.3 cm/s, 0.54-0.64 g) that were proportional to participant height so that perturbations were mechanically similar across participants (Payne et al., 2019; Payne & Ting, 2020). The large perturbation was intentionally designed to be substantially more difficult than typical perturbations, moving backward ∼20 cm in 0.5 s, in order to challenge the young adult population. Platform motion lasted a full duration of 600 ms in all perturbations, which provided similar timing and duration of acceleration, velocity, and displacement across all three perturbation conditions up until the platform decelerated at ∼500 ms (Figure 1).

**Figure 1.**
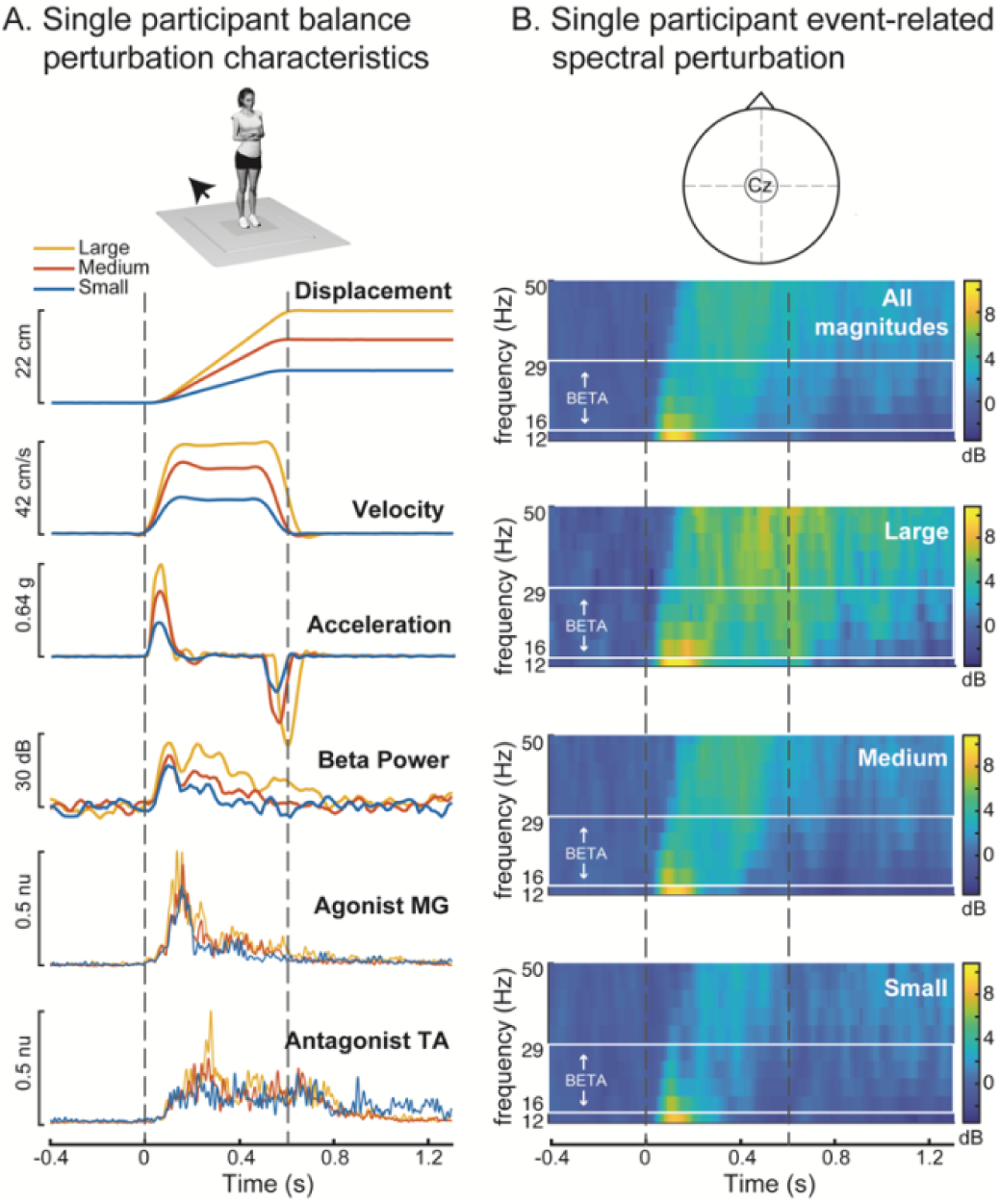
Kinematics of perturbations with evoked cortical and muscle responses shown in an exemplary participant. **A.** Perturbations were scaled in displacement, velocity, and acceleration for each of the 3 conditions. Perturbations for all conditions evoked increases in cortical oscillatory beta (13-30 Hz) power and muscle activity in the medial gastrocnemius (MG) and tibialis anterior (TA). **B.** Event-related spectral perturbation (ERSP) (Cz electrode) across each of the three perturbation conditions in the same exemplar participant. The beta frequency range used for analysis is designated in the solid white box, with the exclusion of lower frequencies enabling higher temporal resolution of beta frequencies. The participant shown in this example (19, F) had relatively low balance ability (0.15), and a prominent second peak of beta activity, which was not observed in all participants. g: units of gravity, dB: decibels, nu: normalized units

The time between each perturbation was at least 14 seconds to ensure that EEG and EMG activity had returned to baseline levels based on visual inspection of real time recordings. As described previously (Payne & Ting, 2020), participants were asked to execute stepping responses on half of perturbations and to resist stepping on the other half, presented in pseudorandom block order. To control for body kinematics and motor control differences within perturbations of the same condition and between participants of different balance abilities, we included only the nonstepping trials in the present analyses to evaluate the cortical contributions to successful feet-in-place reactions. Block randomization was used to balance the presentation of each of the 3 perturbation magnitudes throughout the experiments, with each of eight blocks containing two perturbations of each magnitude in a random order. Each block of six trials was pseudo-randomly assigned one of the instruction sets (i.e. step or keep feet in place), with a maximum of two consecutive blocks before changing the instructions. Across participants, bias due to the order of exposure to perturbation magnitude and instructions were minimized by using three block-randomized orders of the perturbations, and by counterbalancing the instructions across sets (i.e. two versions with opposite instructions)

### Defining successful feet-in-place trials

Platform-mounted force plates (AMTI OR6-6) collected ground reaction forces under each foot during perturbations, which were sampled at 1000 Hz after an online 500 Hz low-pass analog anti-alias filter. Trials in which the participant took an accidental or intentional step were marked by visual inspection in real time and confirmed offline based on the vertical ground reaction forces. Stepping trials were defined as trials in which the vertical load force under either limb reached a value below 5 Newtons within the first 2000ms after perturbation onset and were excluded from subsequent analyses.

### Balance Ability

Balance ability was assessed by balance performance in a challenging beam walking task (Sawers & Ting, 2015) administered after completing the perturbation series. Participants made six attempts to walk across a narrow balance beam (12 feet long, 0.5-inch-wide, 1 inch tall) with their arms crossed in standardized shoes for foot protection. Each trial ended when the participant (1) reached the end of the beam, (2) stepped off the beam, or (3) uncrossed their arms. Balance ability was quantified as the beam distance traversed across six trials and normalized to a maximum possible score of 1 if the participant reached the end of the beam in all 6 trials. A balance ability score close to 0 would indicate the participant consistently lost balance near start of the beam, requiring a recovery step off of the beam.

Participants were not instrumented with EEG, EMG equipment or kinematic markers during the beam walking task.

### Electroencephalography (EEG) data collection and analysis

Brain activity was recorded continuously throughout the balance perturbation protocol using a 32-channel set of actiCAP active recording electrodes (Brain Products GmbH, Germany) placed according to the international 10–20 system, with the exception of electrodes TP9/10 which were placed over the mastoids as reference electrodes. Recordings from the Cz electrode positioned over primary motor and supplementary motor cortical regions were used for analysis. Custom MATLAB scripts and EEGLAB (Delorme & Makeig, 2004) functions were used to epoch (−1000 ms to 2000 ms, relative to perturbation onset at time = 0) and reference EEG activity to the mastoids on a single-trial basis for further analysis.

Time-frequency analyses were performed to assess changes in oscillatory power prior to and during reactive balance recovery. Changes in oscillatory power were quantified in single trial epochs using wavelet time-frequency analyses in EEGLAB (pop_newtimef.m). A tapered Morlet wavelet with three oscillatory cycles at the lowest frequency (11 Hz), linearly increasing up to 6 cycles at the highest frequency (50 Hz) was used to measure power at each frequency in a sliding window of 285 ms. This transformation calculates the event-related spectral perturbation (ERSP), which represents changes in oscillatory power relative to perturbation onset in a defined set of frequencies (Makeig, 1993). ERSP values were calculated for each trial at 200 time points 14 ms apart, and 10 linearly spaced frequencies from 11.7 Hz to 50 Hz based on the spectral and temporal resolution. This resulted in a time course of oscillatory power at a lower sample rate than the input time series data because each measurement of oscillatory power is computed from a window containing many time points of the original time series data. Single-trial ERSP values were averaged across four sampled frequencies within the beta range (centered on 15 Hz, 20 Hz, 24 Hz, and 28 Hz) to obtain a single waveform of the time course of beta frequency power for each trial. Single-trial beta power values were then averaged across trials for each perturbation condition within individuals.

### Electromyography (EMG) data collection and analysis

While primary analyses focused on cortical beta power, we also quantified EMG activity from agonist and antagonist ankle muscles to compare the time course of perturbation-evoked beta power and muscle activity. Surface EMGs (Motion Analysis Systems) were collected bilaterally from primary agonist and antagonist muscles crossing the ankle joint, the medial gastrocnemius muscle (MG) and the tibialis anterior muscle (TA), respectively. Skin was shaved and scrubbed with an isopropyl alcohol wipe before electrode placement using standard procedures (Basmajian & Blumenstein, 1980). Bipolar silver silver-chloride electrodes were used (Nortrode 20, Myotronics, Inc., Kent, WA).

Electromyography signals were sampled at 1000 Hz and anti-alias filtered with an online 500 Hz low-pass filter. Raw EMG signals were epoched between −400 ms and 1400 ms relative to perturbation onset. EMG signals were then high-pass filtered at 35 Hz offline with a third-order zero-lag Butterworth filter, mean-subtracted, half-wave rectified, and subsequently low-pass filtered at 40 Hz. Single-trial EMG data were then normalized to a maximum value of 1 within each participant for left and right sides (e.g. within each participant, left TA activity is normalized such that a maximum value of 1 is observed on at least one trial) and then averaged bilaterally. EMG data was then averaged across trials within each perturbation magnitude within each participant.

#### Quantification of early and late phase evoked beta power within discrete time windows

Pre-perturbation beta activity was quantified as the mean beta power in the 400 ms before perturbation onset. Post-perturbation beta power was initially quantified as the peak beta power in the 400 ms after perturbation onset, prior to the deceleration of the moving platform. To test whether post-perturbation increases in beta power were significant, we performed a two-way ANOVA (factors: time bin and participant) in MATLAB (anova2.m) comparing beta power between these pre- and post-perturbation time bins. The inclusion of the participant factor is necessary to control for between-participant differences when testing the within-participant factor of time bin. Pre-perturbation baseline beta activity was then subtracted from the beta traces for all subsequent analyses. Based on the observation of two clear peaks of beta activity in some individuals (e.g. Figure 1), we defined two additional time bins (50–150 ms and 150-250 ms) of analysis to capture these distinct early and late phase beta power peaks for all participants.

#### Differences in beta power across perturbation magnitude

We compared peak beta power across perturbation magnitudes for each of the post-perturbation time bins (0–400 ms, 50–150 ms, and 150–250 ms) using two-way ANOVAs (factors: perturbation magnitude and participant), using tukey tests (multcompare.m) for post-hoc comparisons across perturbation magnitudes at α = 0.05. The participant factor is included to control for between-participant differences when testing the within-participant factor of perturbation magnitude.

#### Relationship between balance ability and peak beta power

We used generalized linear models (in SAS) to test for associations between peak beta power in each time-bin (0-400 ms, 50-150 ms, and 150–250 ms) and balance ability while accounting for the effect of perturbation magnitude on beta power.

Specifically, the peak beta power amplitudes for each participant and perturbation magnitude were entered into a separate generalized linear model for each time bin, controlling for effects of participant and perturbation magnitude while testing the association to balance ability. We then used univariate linear regressions between balance ability and peak beta power in each time-bin as a post-hoc method to display the associations. The post-hoc regressions display the R2 values from the univariate regression and the p-values obtained from the generalized linear model (which yield the same outcomes as the p-values from the univariate regressions at α = 0.05).

#### Influence of different numbers of nonstepping trials as a potential confound

Some participants had greater difficulty keeping their feet in place in response to the large perturbation, reducing the number of trials available for analysis, which could potentially confound our measures of beta power in a manner related to their balance ability. We chose to use the beam walking test as a measure of balance ability as it has been validated as a sensitive measure of differences in balance ability between healthy young adults with minimal ceiling or floor effects (Sawers & Ting, 2015). In contrast, the percentage of large perturbation trials (N = 8) in which participants were able to resist stepping is a discrete measure, with several participants receiving maximum (always successful) or minimum (never successful) scores.

Nevertheless, there is a possibility that these scores could be associated, which we tested using a univariate linear regression between the balance ability scores and the percentage of trials in which participants successfully maintained their feet in place in the large perturbation, including two individuals who were unable to maintain their feet in place in any trials in the large perturbation. Further, to test whether the number of available trials could have a systematic bias on our beta power measurements, we performed additional linear regressions to test for associations between the percentage of trials in which participants were able to keep their feet in place in the large perturbation and the peak beta power measured across these large perturbations in each of our post-perturbation time bins (0-400 ms, 50-150 ms, 150-250 ms), excluding the two individuals who were unable to maintain feet in place in any large perturbation trials as there were no trials in which to measure their beta power for this analysis.

#### Temporal characterization of evoked cortical and muscle activity using wavelet decomposition

To assess perturbation-evoked changes in cortical oscillatory beta power and evoked muscle activity with greater temporal resolution and statistical power, analyses were performed on wavelet coefficients obtained by an invertible wavelet decomposition. Because wavelet decompositions are more efficient when the number of samples is a power of two (McKay et al., 2013), we shortened our epochs to 128 samples of the beta time course between 400 ms before perturbation onset and ∼1400 ms after perturbation onset prior to wavelet decomposition. Because the statistical power of the wavelet analyses depends on the number of wavelet coefficients, which are determined by the number of samples of the input time-series data (McKay et al., 2013), we additionally down-sampled the EMG data (to 128 samples between −400ms and 1400ms) to match the statistical power between wavelet analyses of cortical beta power and muscle activity. Cortical beta activity and muscle activity were then baseline subtracted again and a discrete wavelet transform was applied to the condition-averaged time-series waveforms (decomposition level 2; wavedec.m) using 3rd order coiflets as the wavelet basis (McKay et al., 2013). This transformed each of the original time-series data waveforms of 128 time points to a wavelet basis containing 160 wavelet coefficients.

### Statistical analyses using wavelet t-tests

To test for significant changes from pre-perturbation baseline in evoked beta or muscle activity at each perturbation magnitude, we performed one-sample t-tests on each of the wavelet coefficients (ttest.m), using a Bonferroni correction to correct for multiple comparisons (yielding α = 0.05/160 = 0.0003). The time course of significant activity was then reverted back to the time-domain by applying the inverse wavelet transform (waverec.m) to a vector containing the mean values of each wavelet coefficient that was significantly greater than zero (p<0.0003125), while the nonsignificant wavelet coefficients were set to zero. Thus, any nonzero activity that remains after the inverse wavelet transform is significantly greater than baseline.

### Identification of activity onset

To compare the time course of significant changes in beta power or muscle activity, we quantified onsets and offsets of the time course of the wavelet reconstructions. Because the wavelet t-tests removed all activity that is not statistically significant, any nonzero activity could be considered as a significant onset. However, the shape of the 3rd order coiflet used in the wavelet decomposition includes small positive ripples to the left and right of each large wavelet peak, which introduces a small nonzero artifact just before the onset of activity. To avoid identifying the artifactual ripple that precedes the onset of activity, we defined nonzero thresholds that were just large enough to pass over these ripples to identify activity onsets. These thresholds were 0.75 dB for beta power and 0.02 normalized units for the muscle activity. Onsets were defined as the first time point at which data exceeded these thresholds after perturbation onset, and offsets were defined as the first subsequent point in which data fell below these thresholds.

### Statistical analyses using wavelet ANOVA

To test for significant differences in evoked beta or muscle activity across perturbation magnitudes, wavelet coefficients were analyzed with a two-way ANOVA (factors: perturbation magnitude and participant) using the methods developed by McKay et al. (2013). Wavelet coefficients corresponding to significant initial F tests (anovan.m) at significance level α = 0.05 were evaluated with post-hoc tukey tests (multcompare.m) relative to the low perturbation magnitude at significance levels Bonferroni-corrected by the number of significant initial F tests (McKay et al., 2013). Statistically significant contrasts in wavelet coefficient magnitude were then transformed back to the time domain (waverec.m), with nonzero values in contrast curves representing the estimated mean of statistically significant differences in beta power or muscle activity in the large or medium perturbations relative to the small perturbation.

### Data availability statement

Single trial EMG and beta time course data will be made publicly available prior to publication. A link to the wavelet ANOVA code will be made available prior to publication. EEGLAB scripts are available at https://sccn.ucsd.edu/eeglab/download.php. Other MATLAB scripts are available through Mathworks (https://www.mathworks.com/) and wavelet analyses require the Signal Processing Toolbox (https://www.mathworks.com/products/signal.html).

## IV. Results

Because participants had greater difficulty responding to the large perturbation without stepping, there were fewer nonstepping trials in the large magnitude to include in our analyses. When asked to recover balance without stepping, participants were able to effectively execute a feet-in-place balance recovery reaction on 98±4% of small perturbations, 96±9% of medium perturbations, and 55±31% of large perturbations. Thus, the average number of trials included for analyses across participants was 8±0 trials for small perturbations, 8±1 trials for medium perturbations, and 4±3 trials for large perturbations. Two participants were unable to recover balance from large perturbations without stepping and were therefore excluded from time bin based comparisons of beta activity across perturbation magnitudes; however, their responses to the small and medium perturbation magnitudes were included in the wavelet analyses.

Differences in the number of available nonstepping trials between participants in the large perturbation were not associated with balance ability scores (F=1.12, p=0.31) or beta power in the large perturbation in any of the post-perturbation time bins (0-400 ms: F=0.00, p=0.95; 50-150 ms: F=0.30, p=0.59; 150-250 ms: F=0.08, p=0.78).

Balance perturbations elicited increases in cortical beta oscillatory power from the baseline mean (67 ± 5 dB) for all conditions, with larger increases in beta power evoked by larger perturbations (Figure 2). In all conditions, an increase in peak beta power was observed during the first 400 ms after perturbation onset (Figure 2A-C, ANOVA, F=44.2, p<0.0001). Following perturbation onset, the magnitude of peak beta power increased with perturbation magnitude in the overall (0-400 ms), early phase (50-150 ms) and late phase (150-250 ms) time bins (Figure 2D, 0-400 ms: F=20.89, p<0.0001, 50-150 ms: F=10.63, p=0.0003, 150-250 ms: F=18.33, p<0.0001). During the overall 0-400 ms time bin, post-hoc tukey tests showed that beta power was significantly greater with each increase in perturbation magnitude (p<0.05). In the early 50-150 ms time bin, beta power was greater in the large compared to the small perturbation (p<0.05), while beta power in the medium perturbation did not differ from the other magnitudes (p>0.05). In the late 150-250 ms time bin, beta power was significantly greater in the large perturbation compared to both the small and medium perturbations (p<0.05) while beta power did not differ between small and medium perturbations (p>0.05)

**Figure 2.**
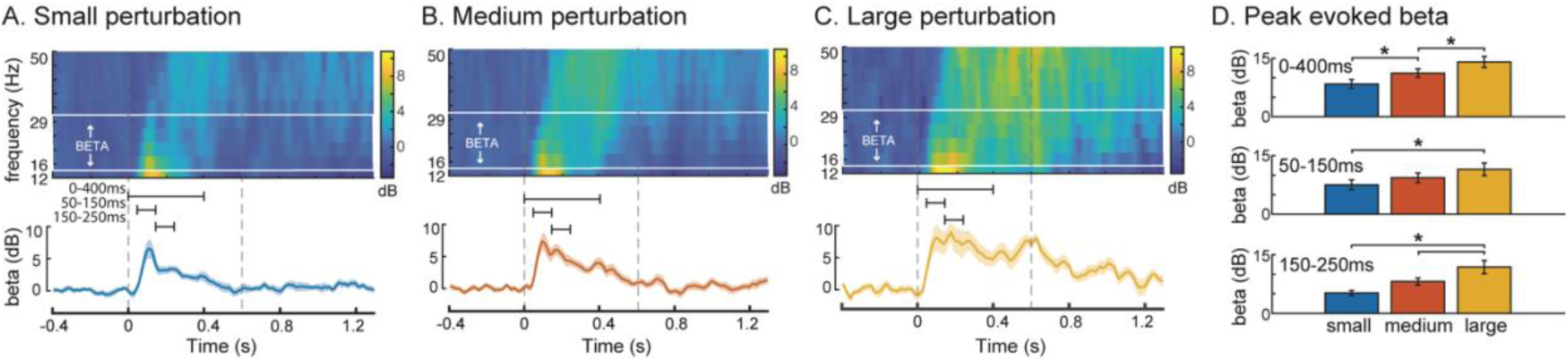
Mean event-related spectral perturbation (ERSP) and beta oscillatory power across participants in (A) small (B) medium and (C) large perturbations. Upper panels show ERSPs, and the solid lines and the shaded regions in the lower panels indicate the mean and standard error of the mean beta power across participants, respectively. Dashed vertical grey lines indicate the perturbation onset and offset. (D) Peak beta power was quantified in each perturbation condition for each of three time-bins indicated by the black horizontal bars in A-C. Asterisks indicate significant differences in beta power between perturbation conditions within each time bin (tukey tests, α = 0.05). dB: decibels

Individuals with lower balance ability had greater perturbation-evoked increases in beta power during the late phase time bin (150-250 ms) of the balance response (Figure 3). Post-perturbation increases in beta power were negatively associated with balance ability in the 150-250 ms time bin (p=0.0037, R2=0.29). In contrast, no association between peak beta power and balance ability was observed during the early phase time bin (50-150 ms, p=0.17) or the overall time bin (0-400 ms, p=0.096).

**Figure 3.**
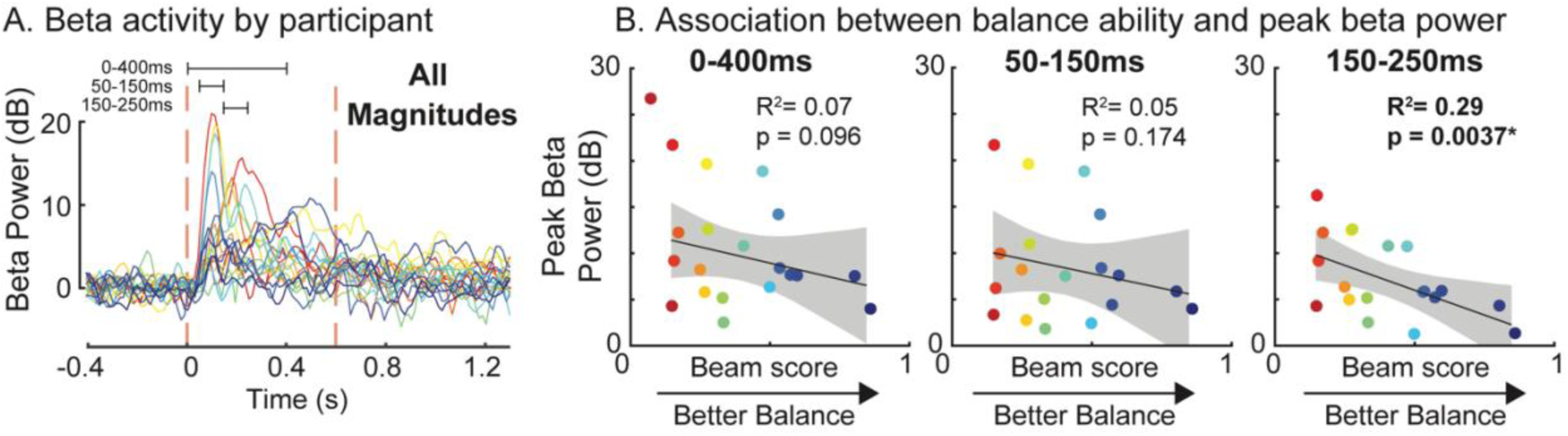
Association between balance ability and individual perturbation-evoked peak beta power in all conditions for each time bin (overall 0-400 ms, early phase 50-150 ms, late phase 150-250 ms). **A.** Time course of mean beta power across all perturbation conditions for each participant. Different colors indicate different participants using a gradient from red (worst balance) to blue (best balance). **B.** Association between peak beta power in each of the three time-bins (0-400 ms, 50-150 ms, 150-250 ms) and balance ability measured as the normalized distance travelled in the challenging beam walking task with values closer to 1 indicating higher balance ability. There was a significant association between balance ability and peak beta power in the 150-250 ms time bin. dB: decibels

Wavelet analyses enabled greater precision in analyzing the time course of changes in beta power relative to muscle activation. In response to balance perturbations in all conditions, beta power increased at approximately 50 ms after perturbation onset across perturbation magnitudes, followed by increases in muscle activity at around 100 ms (Figure 4). Wavelet analyses revealed similar time courses of activity between beta power and muscle activity, with the changes in beta power leading the muscle activity by ∼50 ms, and returning to baseline ∼150-200 ms before the agonist muscle activity (Figure 4AB). Wavelet reconstructions revealed significant increases (wavelet coefficients>0 at p<0.0003) in beta power and muscle activity compared to baseline in all perturbation conditions (Figure 4B). Beta power remained elevated for a longer duration in large perturbations, returning to baseline ∼1150 ms, compared to small and medium perturbations, in which beta power returned to baseline ∼440 ms and ∼550 ms, respectively. In large perturbations both MG and TA activity remained elevated until the end of the window of analysis (>1400 ms). The antagonist TA activity was sustained longer than the agonist MG activity in small (1360 ms vs. 610 ms) and medium (1170 ms vs. 710 ms) perturbations because it was further activated as an agonist by the deceleration of the moving platform (McIlroy & Maki, 1994).

**Figure 4.**
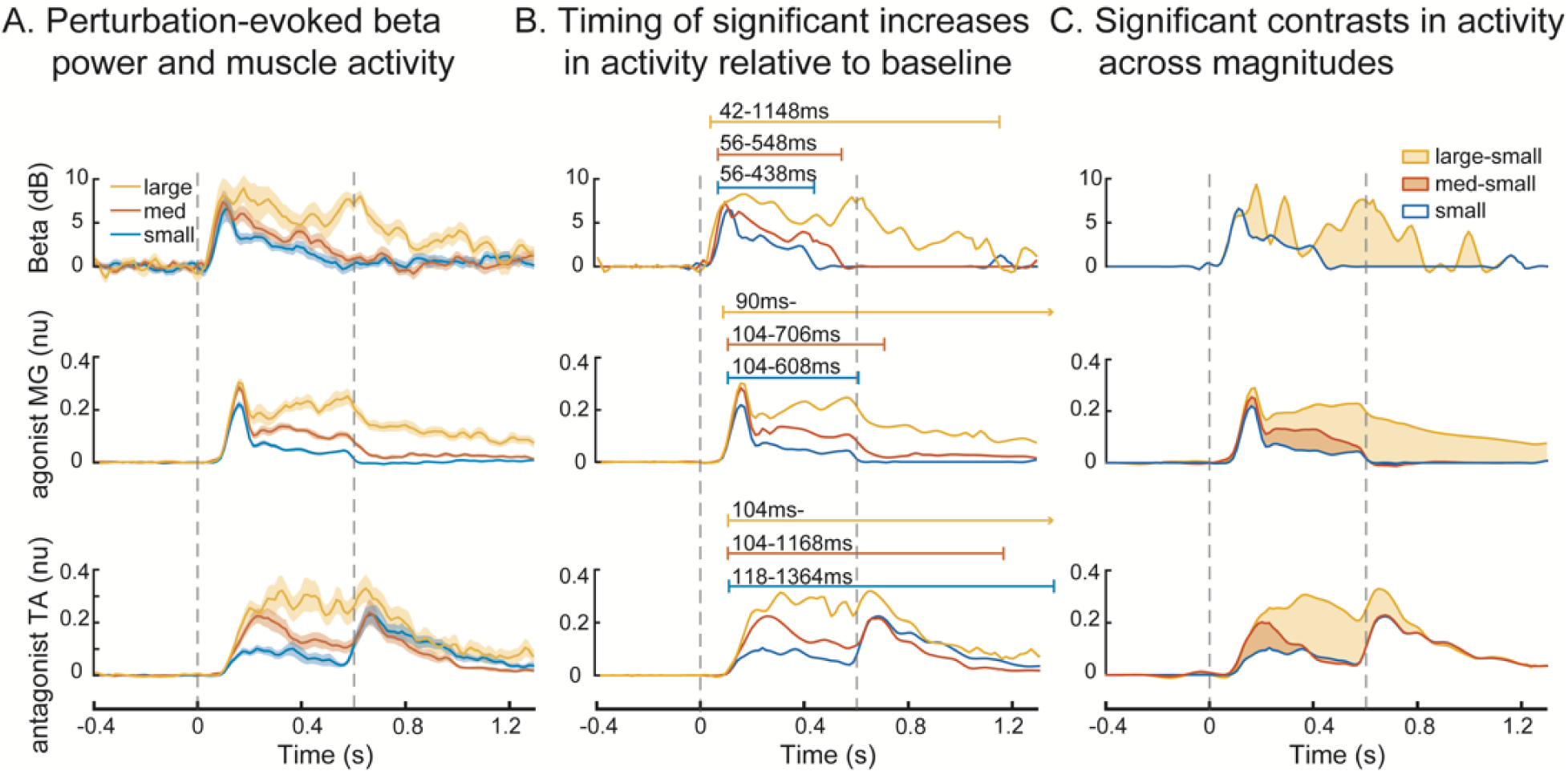
Modulation of cortical beta power and muscle activity across perturbation conditions using wavelet analysis. **A.** Group mean beta power and muscle activity are shown for each perturbation condition, with solid lines showing the group mean and the shaded area showing the standard error of the mean. **B.** Wavelet-reconstructions illustrating the time course of significant changes in evoked beta power and muscle activity relative to baseline. Color-coded horizontal bars indicate time period of elevated activity relative to perturbation onset. **C.** Wavelet contrast curves showing the time course of significant changes (shaded area) in beta power and muscle activity across perturbation conditions relative to the activity in the small perturbation condition. Dashed lines indicate onset and offset of platform movement.

Wavelet contrast curves show differences in the evoked responses across perturbation magnitudes relative to the small perturbation (Figure 4C). Based on the wavelet contrasts, there were no differences in evoked changes in beta power between small and medium perturbations at any time point, and the differences between small and large perturbations began after the initial peak of beta activity (>150 ms), with the greatest difference in beta power between small and large perturbations occurring at the end of platform motion and during continued balance recovery (400-800 ms). In contrast, each increase in perturbation magnitude resulted in an increase in the evoked muscle activity in both agonist and antagonist muscles.

Agonist MG activity was greater in medium vs. small perturbations from the initial peak (∼150 ms) throughout platform motion, while differences in large vs. small perturbations were sustained until the end of the window of analysis throughout balance recovery (>1400 ms). Antagonist TA activity was greater in medium vs. small perturbations from onset (∼100 ms) to ∼300 ms, while differences in large vs. small perturbations were sustained beyond the end of platform motion, ending ∼800 ms.

## V. Discussion

The present study suggests that sensorimotor beta cortical oscillations evoked by balance perturbations provides a neurophysiologic biomarker for cortical engagement in reactive balance recovery. We observed greater modulation of sensorimotor beta oscillations evoked by larger, more challenging balance perturbations, and in individuals with lower balance ability. In contrast to prior studies that have observed beta power modulation beginning prior to the onset of unpredictable balance perturbations (Nakamura et al., 2020; Peterson & Ferris, 2018, 2019; Solis-Escalante et al., 2019; Varghese et al., 2014, 2019), we found modulation of beta power occurred ∼50 ms after the onset of perturbations. Further the time course of evoked beta power was similar to that of evoked balance-correcting muscle activity. As such, perturbation-evoked cortical beta power modulation is not a result of anticipation, but rather may reflect sensory input, sensorimotor integration, and potentially corticomotor engagement during reactive balance recovery. These findings implicate that, in the face of challenging balance conditions, the central nervous system may increase recruitment of cortical resources that engage higher-level sensorimotor integration processes in an effort to maintain upright standing balance. Although the specific functional role of increased cortical beta oscillatory activity remains unclear, our findings identify cortical beta power as a possible biomarker for engagement of sensorimotor cortical brain regions in standing balance control.

### Increased cortical recruitment with increasing balance difficulty

We provide evidence that modulation of cortical beta power occurs in response to balance perturbations and is sustained throughout balance recovery. In contrast to the transient N1 potential that reflects more general neural features of attention (C. E. Little & Woollacott, 2015; Quant et al., 2004) or threat perception (Adkin et al., 2008; Mochizuki et al., 2010), cortical beta power has greater potential for involvement in balance-correcting behavior because it remains elevated throughout balance recovery with a time course comparable to the balance-correcting motor response. Prior studies have shown beta power modulation beginning prior to the onset of unpredictable balance perturbations (Nakamura et al., 2020; Peterson & Ferris, 2018, 2019; Solis-Escalante et al., 2019; Varghese et al., 2014, 2019), making it unclear whether the modulation was occurring in anticipation of, rather than in response to, the balance perturbations. The apparent modulation of beta power prior to perturbation onset in these cases may have resulted from the loss of temporal resolution that occurs with the use of larger sliding time windows that are necessary for the inclusion of lower oscillatory frequencies (e.g. theta) in spectral domain analyses. Focusing specifically on beta power and through the use of wavelet-based functional analyses, we found that beta power modulation occurs in response to balance perturbations at a latency of ∼50 ms. Beta power modulation was sustained throughout platform motion in all perturbation magnitudes and was evoked to a larger extent and longer duration in the large perturbation, persisting throughout balance recovery, which may reflect greater cortical engagement in more difficult perturbations.

Our findings demonstrate that cortical activity is enhanced under more challenging balance conditions and may provide a neurophysiologic biomarker for cortical engagement in balance control. Beta power modulation was larger when participants had greater difficulty recovering balance, either due to more difficult perturbations (within-participants) or due to lower balance ability (between-participants), consistent with our similar findings of the perturbation-evoked cortical N1 response (Payne & Ting, 2020). Under more challenging conditions, the motor demand for balance recovery may exceed the capacity of brainstem-mediated automatic motor responses, necessitating the recruitment of sensorimotor cortical brain regions to recover balance (Jacobs & Horak, 2007). The need for cortical engagement when balance is challenged has been previously inferred on the basis of greater cognitive dual task interference on balance recovery in older adults with balance problems (Shumway-Cook et al., 1997), with such interference effects restricted to later phases (>150 ms) of the muscle activity evoked by perturbations (Rankin et al., 2000). The specific interference on later phases of balance recovery is also in agreement with our finding of an association between balance ability and beta power only in the later (150-250 ms) phase of balance recovery behavior. These results are also consistent with previous research showing brain oscillatory activity in other frequency bands can be modulated as a function of standing postural task challenge, even in the absence of a destabilizing external perturbation (Hülsdünker et al., 2015; Hülsdünker et al., 2016; Sipp et al., 2013). As beta oscillations have also been identified as an early biomarker of motor skill acquisition (Espenhahn et al., 2019), the elevated beta power under more challenging conditions could also index motor skill acquisition or adaptation processes as people are pushed toward their “challenge point” for optimal motor learning (Guadagnoli & Lee, 2004). Though the specific functional role of increased cortical beta oscillatory activity remains unclear, our findings identify cortical beta power as a possible biomarker for engagement of sensorimotor cortical brain regions when balance is challenged.

Although there were fewer nonstepping trials available for some individuals, particularly at higher perturbation magnitudes, differences in the number of trials did not explain our results for within- or between-participant analyses. There was no relationship between nonstepping success rates and individual balance ability in the beam walking task or beta power magnitude, making it unlikely that differences in the available number of trials between participants had a significant impact on our findings. While it may seem surprising that the ability to recover balance without stepping is not associated with our measure of balance ability, prior studies have demonstrated that stepping is often a preferred strategy rather than a point of failure or a “strategy of last resort,” as steps are typically initiated and executed much earlier than necessary (Brian E Maki & McIlroy, 1997). Additionally, nonstepping success rates differ from the continuous measure of balance ability because nonstepping success rates are a discrete measure with few distinct outcomes given so few (N = 8) trials at the most challenging perturbation level, and nonstepping success rates are subject to ceiling and floor effects, with several participants receiving maximum (always successful) or minimum (never successful) outcomes. The association between beta power measured during balance perturbations and a performance measure in a different balance task, i.e., beam walking, suggests these between-participant differences in beta power may reflect cortical engagement related to a more general aspect of balance ability, rather than specific differences in sensory and/or motor activity during the perturbations. People frequently lose and recover their balance during the beam walking task (without stepping off of the beam, e.g. abruptly stopping to bend at the hips or kick a leg out laterally for a few seconds to get their center of mass back over the beam before continuing to walk), consistent with our finding that individuals who are better at reactively recovering balance perform better on the beam walking task. For these reasons, we believe it is meaningful to compare these two balance tasks, and that our outcomes are not confounded by the difficulty some participants had keeping their feet in place in response to the perturbations.

The use of a small number of trials is both a limitation and a strength of the present study. Although the small number of trials limits the precision of our estimate of the average time course of beta oscillatory activity, averaging over a large number of trials may misrepresent the uniformity of both the balance recovery behavior and the beta activity modulations across trials. Balance recovery behavior can adapt rapidly with experience (Horak & Nashner, 1986; Welch & Ting, 2014), with the earliest trials most closely reflecting responses to single events that would occur with falls in real-world situations.

Additionally, a number of recent studies of upper limb behavior have argued that beta oscillations occur as transient events or “bursts,” which only appear to be sustained when looking at averages across trials (Heideman et al., 2020; Jones et al., 2010; S. Little et al., 2019; Shin et al., 2017). These groups have demonstrated that single trial analyses of such beta burst events can predict trial-by-trial differences in perception (Jones et al., 2010; S. Little et al., 2019; Shin et al., 2017) and motor planning (Heideman et al., 2020). The success of single trial analyses in upper limb behaviors, along with the present findings using relatively few trials, suggest the possibility that trial-by-trial variability in beta oscillatory activity may be able to provide insight into trial-by-trial differences in balance recovery behavior in future studies.

### Role of sensorimotor cortical processing in standing balance control

Sensorimotor beta oscillatory activity may initially share an underlying mechanism with the evoked muscle activity. Sensorimotor beta oscillatory activity is sustained throughout balance recovery with a time course resembling balance-correcting muscle activation. Our lab previously demonstrated that during reactive balance recovery the central nervous system directly scales the initial automatic motor response with body motion at a delay of 100 ms (McKay et al., 2020; Welch & Ting, 2008, 2009, 2014). The similarity of the time course of the initial beta activity to that of the motor responses suggests the possibility that a common source of integrated sensory inputs (He et al., 1991; Lockhart & Ting, 2007) may drive both the initial automatic motor response (at ∼100 ms delay) and the early phase of evoked cortical beta activity (at ∼50 ms delay) during balance reactions. However, later phases of the evoked muscle activity (>150 ms) are more variable, and appear to be influenced by cortical activity based on cognitive dual-task interference (Rankin et al., 2000). While the exact role of beta oscillatory activity in balance control remains unclear, it is possible that the later phase (150-250 ms) of beta activity that was associated with balance ability could be influenced by differences in body motion between individuals of different ability levels, and/or this later beta activity could play a role in driving the more variable later phases of the balance-correcting motor responses. While these possibilities remain to be tested in future studies, the finding that sensorimotor beta oscillations are sustained throughout balance recovery and related to balance ability provides mechanistic insight into cortical contributions to balance recovery, which are particularly relevant to populations with impaired balance.

Beta oscillations may also reflect an attempt to rigidly maintain the pre-perturbation upright posture through cortically-driven muscle activation. Beta oscillations are enhanced and sustained during active isometric motor contractions, when a participant rigidly maintains a fixed arm posture (Kilavik et al., 2013; van Wijk et al., 2012). Similar to upper limb studies that have shown greater beta power when maintaining a fixed arm posture compared to when moving the arm to a new posture (Kilavik et al., 2013; van Wijk et al., 2012), Solis-Escalante et al. (2019) recently found beta power was greater when maintaining the feet-in-place posture during balance recovery compared to when recovering balance by taking a step. As such, it is possible that the sustained increase in beta oscillatory power during the most challenging balance conditions reflects greater cortical contributions to muscle activation in attempt to maintain the pre-perturbation posture, which is consistent with our observation of simultaneous activations of agonist and antagonist muscles (i.e. co-contraction). During isometric contractions, cortical beta oscillations are coherent with beta oscillations observed in muscle recordings, which appear to be mediated in part by corticospinal pyramidal tract neurons based on invasive neural recordings in animal models (Baker et al., 1999). Oscillations in the beta frequency range are also consistent with the conduction latencies for signal propagation throughout sensorimotor circuits, including sensorimotor cortical areas, and granger causality analyses have suggested that both afferent and efferent pathways contribute to driving beta oscillations during isometric contractions (Witham et al., 2011). These prior studies support the possibility that sensorimotor beta oscillations during balance recovery may reflect both ascending sensory inputs and cortically-driven motor outputs.

Enhanced cortical beta oscillations during balance recovery may reflect higher level sensorimotor integration and processing involved in balance recovery. Movement-related beta oscillations have been associated with GABAergic network activity (Baker, 2007; Muthukumaraswamy et al., 2013) with primary anatomical origins within thalamocortical sensorimotor networks, including the primary motor cortex (Gaetz & Cheyne, 2006; Jurkiewicz et al., 2006; Salmelin et al., 1995). GABAergic networks play a central role in the modulation of somatosensory processing (Fioravanti et al., 2019) and can influence corticomotor output through inhibitory and reciprocal connections with pyramidal cells (Yamawaki et al., 2008). A relationship between beta oscillations and somatosensory processing is also supported by larger somatosensory-evoked potential amplitudes when evoked during spontaneous phasic increases in beta power (Lalo et al., 2007). In contrast to volitional movement, where disinhibition of corticomotor output is necessary for movement initiation (Murase et al., 2004; Pfurtscheller & Lopes da Silva, 1999), reactive balance control may demand greater engagement of GABAergic networks for rapid processing of somatosensory information and subsequent modulation of corticospinal output for later phase (>150 ms) balance reactions (i.e. enhanced cortical beta activity). As a next step, studies may investigate how the therapeutic modulation of cortical beta oscillations through pharmacological agents or noninvasive brain stimulation influences reactive balance control. Building upon findings of the present study, future research could provide a clearer picture of the complex relationship between cortical beta oscillations, cortical inhibition and excitation, and human balance control.

### Conclusions

The engagement of sensorimotor cortical brain regions is increased under challenging balance conditions in healthy young adults. Perturbation-evoked cortical beta power could provide a useful biomarker of sensorimotor engagement reflecting an individual’s balance ability. These findings have important implications for the study of balance-correcting behavior in older adults and balance-impaired neurologic patient populations, where the cerebral cortex may have a greater role in balance and postural control.

## Acknowledgments

This research was supported by the Eunice Kennedy Shriver National Institutes of Child Health & Human Development of the National Institutes of Health [R01 HD46922-10, F32HD096816, K12HD055931], the National Institute of Neurological Disorders and Stroke of the National Institutes of Health [1P50NS098685], the National Institute on Drug Abuse of the National Institutes of Health [5T90DA032466], the Division of Emerging Frontiers and Multidisciplinary Activities at the National Science Foundation (1137229), the Georgia Tech Neural Engineering Center, the President’s Undergraduate Research Award provided by the Undergraduate Research Opportunities Program at Georgia Tech, and the Andy Zebrowitz Memorial Brain Research Fellowship Award (2017-2018).

